# *Areca catechu*-(Betel-nut)-induced whole transcriptome changes associated with diabetes, obesity and metabolic syndrome in a human monocyte cell line

**DOI:** 10.1101/2020.08.03.233932

**Authors:** Shirleny Cardoso, B. William Ogunkolade, Rob Lowe, Emanuel Savage, Charles A Mein, Barbara J Boucher, Graham A Hitman

## Abstract

Betel-nut consumption is the fourth most common addictive habit globally and there is good evidence to link it with obesity, type 2 diabetes and the metabolic syndrome. We adopted a genome-wide transcriptomic approach in a human monocyte cell line incubated with arecoline and its nitrosated products to identify gene expression changes relevant to obesity, type 2 diabetes and the metabolic syndrome. The THP1 monocyte cell line was incubated separately with arecoline and 3-methylnitrosaminopropionaldehyde (MNPA) in triplicate for 24 hours and pooled cDNA indexed paired-end libraries were sequenced (Illumina NextSeq 500). After incubation with arecoline and MNPA, 15 and 39 genes respectively had significant changes in their expression (q<0.05, log fold change 1.5). Eighteen of those genes have reported associations with type 2 diabetes and obesity in humans; of these genes there was strong evidence to implicate *CLEC10A*, *MAPK8IP1*, *NEGR1*, *NQ01* and *INHBE*. In summary, these pilot studies have identified a large number of genes whose expression was changed significantly in human TPH1 cells following incubation with arecoline or with 3-methylnitrosaminopropionaldehyde. These findings suggest that further investigation of these genes in betel-quid chewers with obesity and/or type 2 diabetes is warranted.

## Introduction

Obesity and Type 2 diabetes are reaching epidemic proportions worldwide, but particularly so in South Asian communities living in the Indian-subcontinent or who have migrated to other countries[1]. In the UK there is a 3 to 4 fold increase in type 2 diabetes prevalence in South Asians compared to the general population; furthermore the disease manifests at a 10-15 years younger age and is strongly associated with the metabolic syndrome and cardiovascular disease. Apart from lifestyle, potentially reversible environmental factors driving this disease are largely unknown.

Betel quid consumption is the fourth most common addictive habit, used by 10% of the global population and very common in South Asians. The link between cancer risks (oropharyngeal, oesophageal and hepatocellular) and the ‘betel-chewing’ habit is well established[2–4]. Evidence also implicates an association between betel consumption and obesity, the metabolic syndrome and type 2 diabetes. In a meta-analysis of 17 Asian studies, betel quid chewing was found to be significantly associated with obesity (relative risk (RR) = 1.47), metabolic syndrome (RR=1.51), diabetes (RR=1.47), hypertension (RR=1.45) and cardiovascular disease (cardiovascular disease: RR=1.2)[5]. Furthermore, in the Keelung Community Integrated Screening programme studies from Taiwan, paternal betel-chewing was associated, dose-wise, with increases in early onset metabolic syndrome in their never-chewing offspring, while betel-chewing in adults increased their risks, dose-wise, of early onset type 2 diabetes and cardiovascular disease [6–8]. These data in humans support earlier data reported in CD1 mice, where it was found that a proportion of betel-fed adult mice developed hyperglycaemia and obesity and, more remarkably, that amongst their never-betel-fed offspring hyperglycaemia was detected in up to the 4^th^ generation and that the vertical transmission of hyperglycaemia was associated with paternal, but not maternal, hyperglycaemia[9].

The betel quids (also known as paan) are usually contain betel (areca) nut, slaked lime and sometimes tobacco wrapped in betel leaf[10]. In Taiwan tobacco is not used. Nitrosation of the major arecal alkaloid, arecoline, forms 3-methylnitrosaminopropionaldehyde (MNPA), and 3-methylnitrosaminopropionitrile (MNPN) and both these compounds have been identified as carcinogens [11]. Many nitroso-compounds (e.g. streptozotocin) have been reported as being diabetogenic, low doses causing type 2 diabetes -like diabetes, suggesting that betel-chewing might be one of the aetiological factors for the increases in type 2 diabetes and associated metabolic disorders in South Asians[12, 13]. Mechanisms that might link betel chewing with these disorders include inflammation, increases in hepatic synthesis of lipids and glucose, in adipogenesis, in adipose tissue glucose uptake, reductions in lipolysis and glycolysis, neurological, hepatic or intestinal effects on appetite and adverse effects on vitamin D metabolism[14, 15].

In the present proof of principle study we sought to investigate possible biological links between type 2 diabetes (and related disorders) and exposure to arecoline and its nitrosated products in a human monocyte derived cell line (THP1) using a whole transcriptome sequencing approach. THP1 was chosen due the central role that low-grade inflammation plays in the underlying causes and progression of obesity (for instance adipose tissue macrophages), metabolic syndrome and related disorders including both cardiovascular disease and type 2 diabetes[16, 17].

## Materials and Methods

The THP1 (human acute monocytic leukemia derived; ATCC number TIB-202 purchased from Thermofisher) cell line[18] was regularly maintained in RPMI 1640 medium containing GlutaMAX™, supplemented with 10% FCS (Gibco™ Newborn Calf Serum [heat inactivated], of New Zealand origin; Thermo Fisher Scientific), 5% AA (Gibco® MEM), Non-Essential Amino Acids, 100 U/mL penicillin and 100 μg/mL streptomycin (Thermo Fisher Scientific). Cells were grown at 37 °C in a humidified atmosphere of 5% CO2 in air, and sub-passaged with fresh complete RPMI medium every three days. The cell line was regularly checked to be mycoplasma free using the VenorGeM Mycoplasma detection kit (Cambio Ltd, UK) according to manufacturer’s instructions.

For chemical treatments, arecoline, MNPA and PMA (Phorbol-12-Myrsitate-13-Acetate) were diluted in 100 % methanol while MNPN was diluted in 100 % ethyl acetate. 1×10^6^ THP1 cells were treated with either 100 ng/ml Arecoline, 2-5 ng/ml MNPA, 50 ng/ml MNPN or 200 ng/ml PMA as a positive control in 6-well plates and cells were harvested after 6 h, 24 h and 48 h of treatment. Methanol and ethyl acetate were used as negative controls. Three independent experiments were performed for each exposure. All chemicals were purchased from Sigma-Aldrich.

### RNA extraction and RT-qPCR for gene expression of human TNFa, IL-6 and IL-8 analysis

Total RNA was extracted from treated cells using QIAGEN RNeasy Kit according to manufacturer’s instructions. cDNA was synthesized using 1 μg of the extracted RNA with an Oligo (dT) primer using a SuperScript® IV First-Strand Synthesis System (Thermo Fisher Scientific) as follows: primer annealing at 65 °C for 5 min; RNA reverse transcriptase at 50 °C for 1 h 10 min and at 70 °C for 15 min. The cDNA was used as a template to determine the expression level of human TNFa, IL-6, IL-8 and 18S in (arecoline, MNPA or MNPN) treated/untreated THP1 cells. The RT-PCR was performed on StepOne™ Real-Time PCR System thermal cycler (Applied Biosystems™). Each PCR reaction consisted of 2 μl of cDNA, 2X SYBR® Green JumpStart™ Taq ReadyMix™ (Sigma-Aldrich), 0.2 μM of each forward and reverse primers (table S1). qPCR reaction conditions were: cDNA denaturation at 95 °C for 5 min, cDNA amplification at 95 °C for 15 sec, primer annealing at 62 °C for 1 min for 45 cycles, then melt curves were obtained at 95 °C for 15 s, 60 °C for 1 min and a final step at 95 °C for 15 s. All target genes were normalised to *18S RNA* using the standard *ΔΔ*Ct method. Results were analysed with Thermo Fisher StepOne software v2.3. Each experiment was performed in duplicate and fold change expression level was reported relative to 18S level.

### RNA-sequencing and bioinformatics

Fragmented cDNA Sequencing libraries were generated from 100ng of Total RNA using NEBNext Ultra II with polyA isolation module (Illumina, San Diego, California, USA) according to manufacturer’s protocol. cDNA quantity and quality were evaluated using the Qubit dsDNA HS assay kit (Thermo Fisher Scientific, Erembodegem-Aalst, Belgium). Size distribution of our library was determined using an Agilent 2100 Bioanalyser. Pooled indexed paired-end libraries were sequenced on the Illumina NextSeq 500 (Illumina, USA) using the manufacturer’s instructions. Sequencing was performed at Queen Mary University of London Genome Centre core facility in the Blizard Institute London.

Sequenced reads were mapped using Kallisto[19] with default settings. Mean insert sizes and standard deviations were provided as input. Analysis of differential gene expression was performed using sleuth, applying a generalized linear model and utilising bootstraps on reads to estimate inferential variance. Genome-wide corrected p-values were calculated using the Bonferroni multiple testing adjustment procedure.

Functional annotation as well as pathway enrichment analyses were performed using DAVID, Reactome and Metascape ((https://metascape.org/gp/index.html#/main/step1).

### Candidacy of genes identified were assessed by look-ups in

1. GeneCards: The human genome database (https://www.genecards.org/) to check for alias’s, gene summary (Entrez, Genecards and UniProtKB/Swiss-Prot) and mRNA expression in normal human tissues (GTEx, Illumina, BioGPS)
2. Type 2 diabetes knowledge portal (http://www.type2diabetesgenetics.org/). Genes considered were only those with strong evidence for signal defined by either at least one variant within the coding sequence ± 100kb that is associated with at least one phenotype with a p value<5e-8 identified by a genome-wide association scan (GWAS), or at least one known variant with a missense or protein truncating mutation in the encoded protein that is associated with at least one phenotype with a p value <5e-6.
3. PubMed (GAH and BJB independently) by searching biology of the identified gene and biological relevance to obesity, type 2 diabetes or the metabolic syndrome before collation of results.

## Results

### Baseline experiments in triplicate

Whilst an inflammatory response was seen after incubation with either arecoline, MNPA or PMA, no response was found at 6, 24 or 48 hours with MNPN – it was therefore decided that we could not proceed further with investigation of the last of these compounds.

After 24 hours incubation with PMA increased expression was found for TNFA, IL6 and IL8 (mean fold change 2.5, 20.6 and 23.6 units respectively); arecoline incubation increased IL6 expression (mean fold change 2.8) and MNPA incubation also increased IL6 expression (mean fold change 2.8). PMA increased expression at 48 hours for IL6 and IL8. We therefore decided to proceed with the incubations at 24 hours and whole transcriptome expression experiments were run in triplicate.

### Whole transcriptome analysis

#### Incubation with arecoline

275 gene hits were identified with a q<0.05 (table S2) reducing to 15 with a log-fold change in either direction of 1.5 (table 1). Amongst the 15 genes, 4 genes have relevance to diabetes, obesity and/or metabolic syndrome listed below

**Table 1.** Genes identified after incubation with arecoline

| Gene Name | Gene ID | Gene Name | q-value | logfc | Gene Description |
| --- | --- | --- | --- | --- | --- |
| H3F3AP4 | ENSG00000235655 | H3F3AP4 | 0.0328274 | -6.7682546 | H3 Histone Pseudogene 6 |
| MEP1A | ENSG00000112818 | MEP1A | 0.0432902 | -2.1619323 | Meprin A Subunit Alpha |
| KBTBD11-OT1 | ENSG00000283239 | KBTBD11-OT1 | 0.0461988 | -3.7981785 | KBTBD11 Overlapping Transcript 1 |
| IGFBP3 | ENSG00000146674 | IGFBP3 | 0.0432902 | -2.2897729 | Insulin Like Growth Factor Binding Protein 3 |
| AC097372.1 | ENSG00000250673 | AC097372.1 | 0.0435549 | -2.1772391 | Reeler Domain Containing 1 |
| PTCRA | ENSG00000171611 | PTCRA | 0.0432902 | -1.9842384 | Pre T Cell Antigen Receptor Alpha |
| IL3RA | ENSG00000185291 | IL3RA | 0.0420978 | -1.9557171 | Interleukin 3 Receptor Subunit Alpha |
| CLEC10A | ENSG00000132514 | CLEC10A | 0.0332128 | -1.8776835 | C-Type Lectin Domain Containing 10A |
| ACKR3 | ENSG00000144476 | ACKR3 | 0.0432902 | -1.7457949 | Atypical Chemokine Receptor 3 |
| JUP | ENSG00000173801 | JUP | 0.0354724 | -1.5030002 | Junction Plakoglobin |
| MAPK8IP1 | ENSG00000121653 | MAPK8IP1 | 0.0461988 | -1.5085173 | Mitogen-Activated Protein Kinase 8 Interacting Protein 1* |
| MATK | ENSG00000007264 | MATK | 0.0295919 | -1.5132972 | Megakaryocyte-Associated Tyrosine Kinase |
| TREML3P | ENSG00000184106 | TREML3P | 0.0328274 | 1.93092581 | Triggering Receptor Expressed On Myeloid Cells Like 3, Pseudogene |
| AC245036.5 | ENSG00000269271 | AC245036.5 | 0.0432902 | 2.98837483 | RNA gene; lncRNA |
| AC113189.4 | ENSG00000272884 | AC113189.4 | 0.0466794 | 2.75155456 | RNA gene; lncRNA |

Listed genes satisfied the following criteria $q < 0.05$ and a log-fold change of 1.5

a. Insulin Like Growth Factor Binding Protein 3 (*IGFB3* non-logged fold change 0.08),
b. C-Type Lectin Domain Containing 10A (*CLEC10A* fold change 0.14),
c. Junction Plakoglobin (*JUP* fold change 0.21)
d. Mitogen-Activated Protein Kinase 8 Interacting Protein 1 (*MAPK8IP1* fold change 0.21)

Pathway (Metascape) analysis of the 275 genes (figure 1) listed in the online table 2, revealed 5 significant pathways after statistical correction: myeloid cell activation involved in immune response, cellular response to thyroid hormone stimulus, responses to toxic substances and Hematopoietic cell lineage.

**Figure 1.**
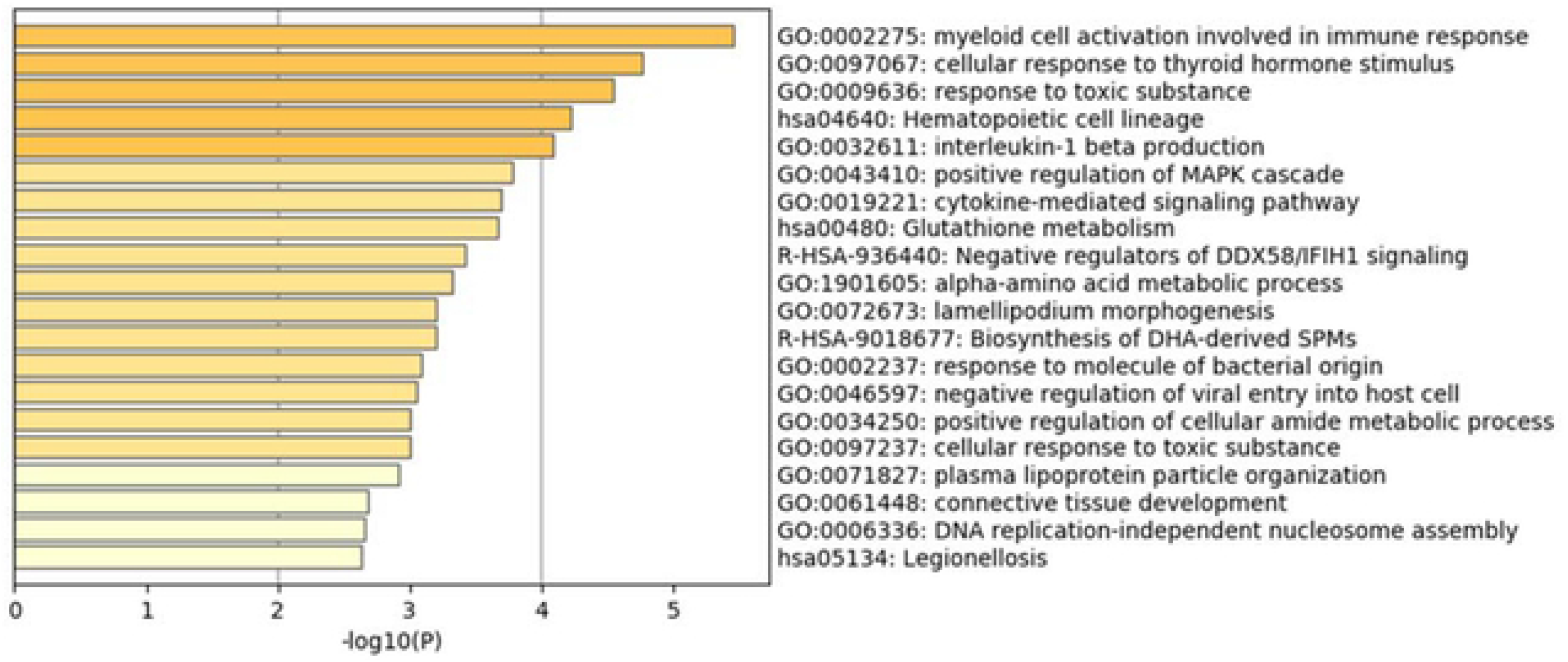
Pathway (Metascape) analysis arecoline incubation. Metascape Bar plot (P value (log10 scale)) showing Top 20 arecoline-induced, enriched functional ontology clusters (GO and KEGG terms) one per cluster.

**Table 2.**
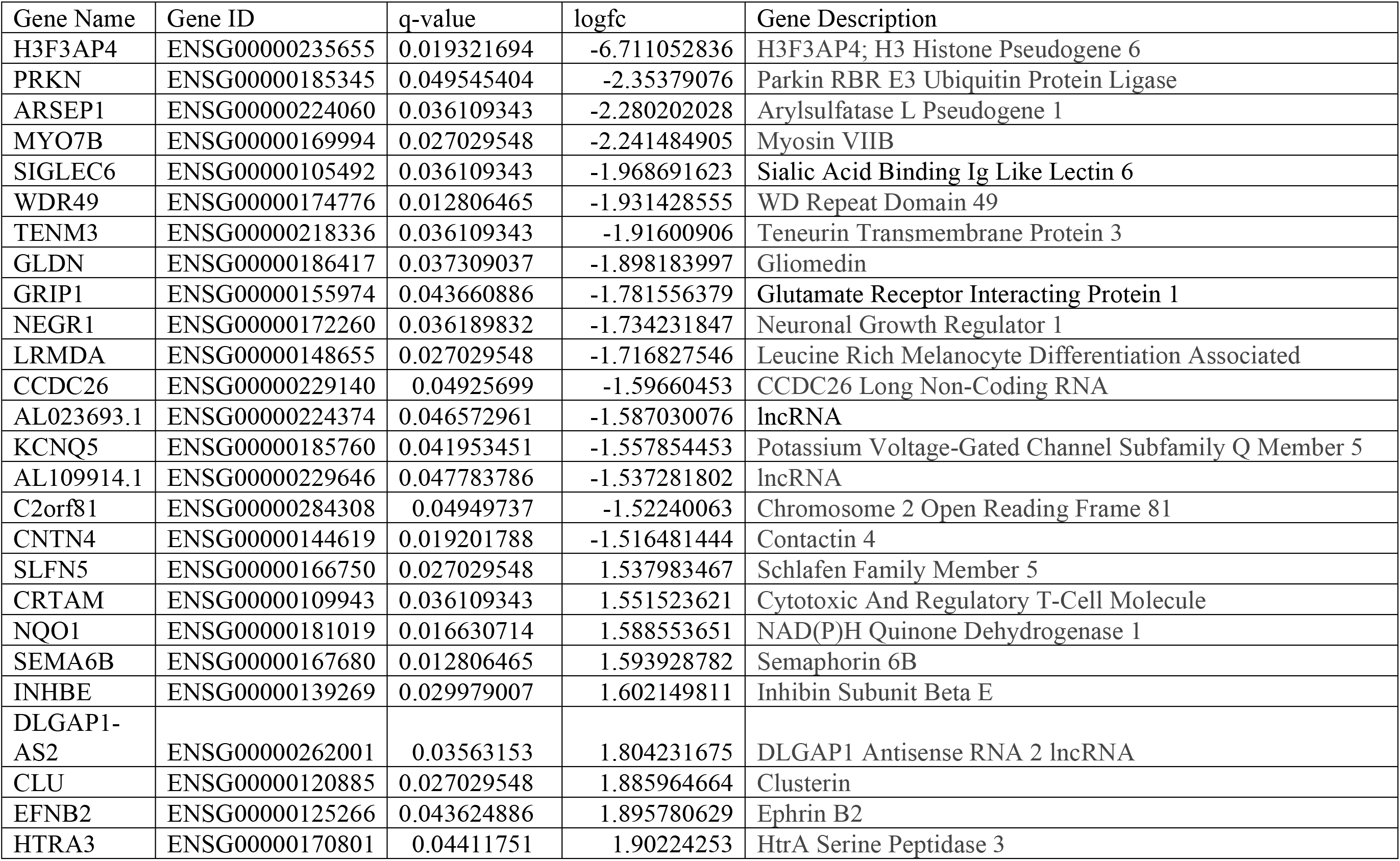
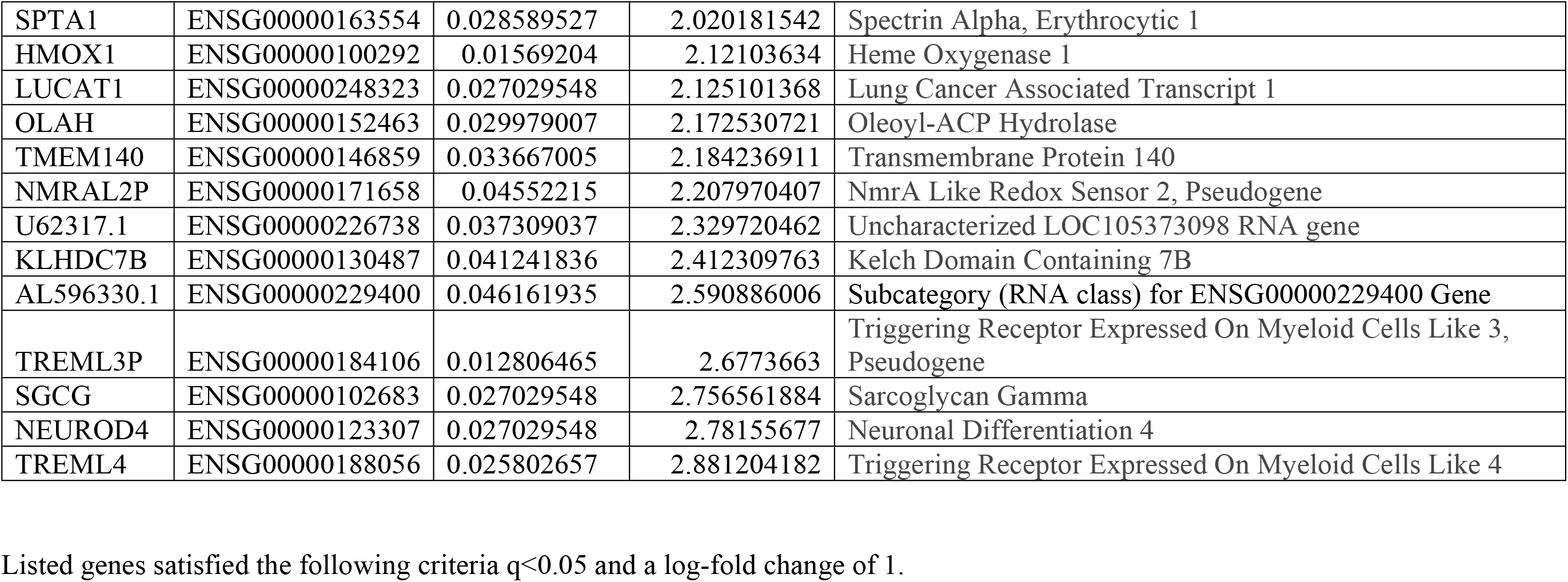
Genes identified after incubation with MNPA

| Gene Name | Gene ID | q-value | logfc | Gene Description |
| --- | --- | --- | --- | --- |
| H3F3AP4 | ENSG00000235655 | 0.019321694 | -6.711052836 | H3F3AP4; H3 Histone Pseudogene 6 |
| PRKN | ENSG00000185345 | 0.049545404 | -2.35379076 | Parkin RBR E3 Ubiquitin Protein Ligase |
| ARSEP1 | ENSG00000224060 | 0.036109343 | -2.280202028 | Arylsulfatase L Pseudogene 1 |
| MYO7B | ENSG00000169994 | 0.027029548 | -2.241484905 | Myosin VIIB |
| SIGLEC6 | ENSG00000105492 | 0.036109343 | -1.968691623 | Sialic Acid Binding Ig Like Lectin 6 |
| WDR49 | ENSG00000174776 | 0.012806465 | -1.931428555 | WD Repeat Domain 49 |
| TENM3 | ENSG00000218336 | 0.036109343 | -1.91600906 | Teneurin Transmembrane Protein 3 |
| GLDN | ENSG00000186417 | 0.037309037 | -1.898183997 | Gliomedin |
| GRIP1 | ENSG00000155974 | 0.043660886 | -1.781556379 | Glutamate Receptor Interacting Protein 1 |
| NEGR1 | ENSG00000172260 | 0.036189832 | -1.734231847 | Neuronal Growth Regulator 1 |
| LRMDA | ENSG00000148655 | 0.027029548 | -1.716827546 | Leucine Rich Melanocyte Differentiation Associated |
| CCDC26 | ENSG00000229140 | 0.04925699 | -1.59660453 | CCDC26 Long Non-Coding RNA |
| AL023693.1 | ENSG00000224374 | 0.046572961 | -1.587030076 | lncRNA |
| KCNQ5 | ENSG00000185760 | 0.041953451 | -1.557854453 | Potassium Voltage-Gated Channel Subfamily Q Member 5 |
| AL109914.1 | ENSG00000229646 | 0.047783786 | -1.537281802 | lncRNA |
| C2orf81 | ENSG00000284308 | 0.04949737 | -1.52240063 | Chromosome 2 Open Reading Frame 81 |
| CNTN4 | ENSG00000144619 | 0.019201788 | -1.516481444 | Contactin 4 |
| SLFN5 | ENSG00000166750 | 0.027029548 | 1.537983467 | Schlafen Family Member 5 |
| CRTAM | ENSG00000109943 | 0.036109343 | 1.551523621 | Cytotoxic And Regulatory T-Cell Molecule |
| NQO1 | ENSG00000181019 | 0.016630714 | 1.588553651 | NAD(P)H Quinone Dehydrogenase 1 |
| SEMA6B | ENSG00000167680 | 0.012806465 | 1.593928782 | Semaphorin 6B |
| INHBE | ENSG00000139269 | 0.029979007 | 1.602149811 | Inhibin Subunit Beta E |
| DLGAP1-AS2 | ENSG00000262001 | 0.03563153 | 1.804231675 | DLGAP1 Antisense RNA 2 lncRNA |
| CLU | ENSG00000120885 | 0.027029548 | 1.885964664 | Clusterin |
| EFNB2 | ENSG00000125266 | 0.043624886 | 1.895780629 | Ephrin B2 |
| HTRA3 | ENSG00000170801 | 0.04411751 | 1.90224253 | HtrA Serine Peptidase 3 |
| SPTA1 | ENSG00000163554 | 0.028589527 | 2.020181542 | Spectrin Alpha, Erythrocytic 1 |
| HMOX1 | ENSG00000100292 | 0.01569204 | 2.12103634 | Heme Oxygenase 1 |
| LUCAT1 | ENSG00000248323 | 0.027029548 | 2.125101368 | Lung Cancer Associated Transcript 1 |
| OLAH | ENSG00000152463 | 0.029979007 | 2.172530721 | Oleoyl-ACP Hydrolase |
| TMEM140 | ENSG00000146859 | 0.033667005 | 2.184236911 | Transmembrane Protein 140 |
| NMRAL2P | ENSG00000171658 | 0.04552215 | 2.207970407 | NmrA Like Redox Sensor 2, Pseudogene |
| U62317.1 | ENSG00000226738 | 0.037309037 | 2.329720462 | Uncharacterized LOC105373098 RNA gene |
| KLHDC7B | ENSG00000130487 | 0.041241836 | 2.412309763 | Kelch Domain Containing 7B |
| AL596330.1 | ENSG00000229400 | 0.046161935 | 2.590886006 | Subcategory (RNA class) for ENSG00000229400 Gene |
| TREML3P | ENSG00000184106 | 0.012806465 | 2.6773663 | Triggering Receptor Expressed On Myeloid Cells Like 3, Pseudogene |
| SGCG | ENSG00000102683 | 0.027029548 | 2.756561884 | Sarcoglycan Gamma |
| NEUROD4 | ENSG00000123307 | 0.027029548 | 2.78155677 | Neuronal Differentiation 4 |
| TREML4 | ENSG00000188056 | 0.025802657 | 2.881204182 | Triggering Receptor Expressed On Myeloid Cells Like 4 |

Listed genes satisfied the following criteria $q < 0.05$ and a log-fold change of 1.

#### Incubation with MNPA

359 gene hits were identified after incubation with MNPA with a q<0.05 (table S3) reducing to 39 with a log-fold change of +/−1.5 (table 2). Amongst the 39 genes, 14 have relevance to diabetes, obesity and/or metabolic syndrome listed below

a. Gliomedin (*GLDN* fold change 0.11)
b. Glutamate Receptor Interacting Protein 1 (*GRIP1* fold change 0.15)
c. Neuronal Growth Regulator 1 (*NEGR1* fold change 0.14)
d. Potassium Voltage-Gated Channel Subfamily Q Member 5 (*KCNQ5* fold change 0.21)
e. Cytotoxic And Regulatory T-Cell Molecule (*CRTAM* fold change 4.9)
f. NAD(P)H Quinone Dehydrogenase 1 (*NQO1* fold change 4.9)
g. Semaphorin 6B (*SEMA6B* fold change 4.9)
h. Inhibin Subunit Beta E (*INHBE* fold change 5.4)
i. Clusterin (*CLU* fold change 6.7)
j. Spectrin Alpha, Erythrocytic 1 (*SPTA1* fold change 9.3)
k. Heme Oxygenase 1 (*HMOX1* fold change 8.4)
l. Transmembrane Protein 140 (*TMEM140* fold change 10.0)
m. Sarcoglycan Gamma (*SGCG* fold change 24.8)
n. Triggering Receptor Expressed On Myeloid Cells Like 4 (*TREML4* fold change 26.7)

Pathway (Metascape) analysis of the 359 genes (figure 2) listed in the online table 2, revealed 5 significant pathways after statistical correction: regulation of cell adhesion, response to inorganic substances, apoptotic signalling pathway, response to toxic substances and regulation of the innate immune response.

**Figure 2.**
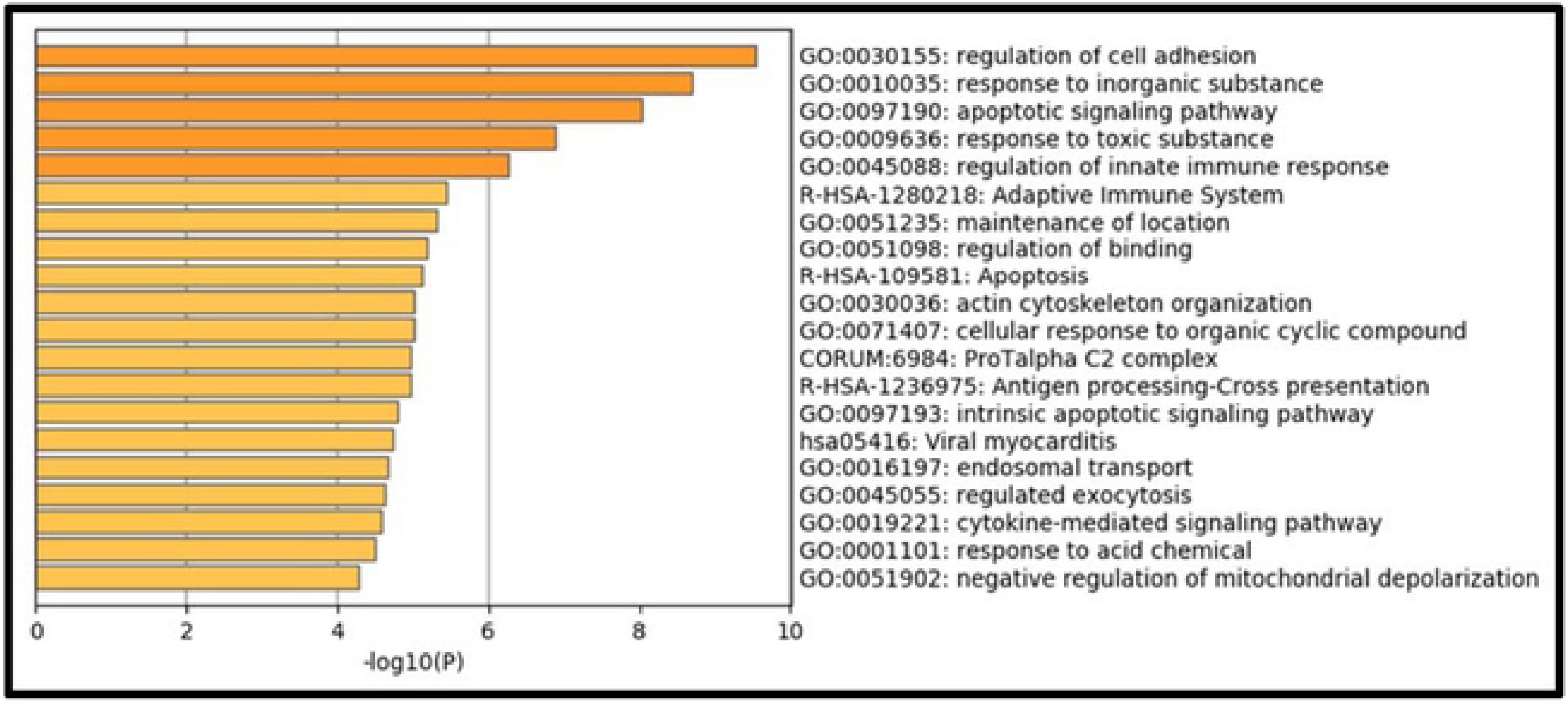
Pathway (Metascape) analysis MNPA incubation. Metascape Bar plot (P value (log10 scale)) showing Top 20 MNPA-induced, enriched functional ontology clusters (GO and KEGG terms) one per cluster.

## Discussion

Whole transcriptome analysis by RNASeq of the human monocyte line THP1 reveals a significant number of genes that are either downregulated or upregulated in response to incubation with arecoline or with MNPA. The aim of our study was to identify genes associated with diabetes, obesity and metabolic syndrome whose expression was significantly altered by exposure to the arecal compounds arecoline and its nitrosated metabolite, 3-methylnitrosaminopropionaldehyde (MNPA). It was also hoped to determine whether the strength of the evidence for any of those genes significantly affected might warrant further investigation amongst betel-chewing communities with a high prevalence of metabolic syndrome related disorders including obesity and type 2 diabetes.

Consistent with the established effects of betel nut ingestion in humans, a number of cellular pathways and genes have been identified as being significantly affected by the arecal compounds used in our approach; these genes are known to relate to immune responses, to cell differentiation and lineage, to responses to toxic and inorganic substances and to the development of obesity and type 2 diabetes in humans. Other genes with significantly altered expression are known to be associated with carcinogenesis and immune function and a smaller number of affected genes are related to neural development and could be associated with addiction.

### Genes of relevance to obesity, diabetes and metabolic syndrome with good evidence that arecoline and/or MNPA may alter its expression based on published studies and genetic evidence include

**C-Type Lectin Domain Containing 10A (*CLEC10A*)** is a calcium dependent endocytic receptor also known as the macrophage galactose-type lectin (MGL or CD301). It has been demonstrated to have a role in regulating adaptive and innate immune responses and is expressed in adipose tissue macrophages where it is associated with phenotypic switching of ATM subclasses in mice that then demonstrate either a lean or an obese phenotype[20]. Furthermore, evidence in humans demonstrates that missense and protein truncating mutations of the CLEC10A gene are strongly associated with the development of type 2 diabetes (http://www.type2diabetesgenetics.org/gene/geneInfo/CLEC10A). In earlier rodent experiments MGL1 was found to be a novel regulator of monocyte trafficking in adipose tissue in response to dietary induced obesity[20, 21].

**Mitogen-Activated Protein Kinase 8 Interacting Protein 1 (*MAPK8IP1*)** gene encodes a regulator of pancreatic beta-cell function; it is also expressed in a large number of tissues including many associated with immune function and is also a trans-activator of the glucose transporter GLUT2. *MAPK8IP1* has a strong association with type 2 diabetes with a missense mutation found in one family and, in *vitro,* that mutation was found to be a key down-regulator of beta cell function[22].

**Neuronal Growth Regulator 1 (*NEGR1*)** is involved in cell adhesion and certain mutations of this gene lead to Niemannn-Pick disease, a rare inherited metabolic disorder. Multiple genome-wide association studies demonstrate strong genome-wide association (GWAS) signals for this gene with BMI, waist circumference and type 2 diabetes[23] (http://www.type2diabetesgenetics.org/gene/geneInfo/NEGR1). Furthermore, *NEGR1* knockout mice develop increased adiposity including increased hepatocyte fat deposition together with increases in glycaemia and in fasting serum insulin levels[24].

**NAD(P)H Quinone Dehydrogenase 1 (*NQO1*)** gene is a member of the NAD(P)H dehydrogenase (quinone) family and encodes a cytoplasmic 2-electron reductase (Entrez Gene) and is part of the antioxidant defence system. There is strong genetic evidence to support an association between *NQO1* variants by GWAS and increased risks of type 2 diabetes and increased BMI (http://www.type2diabetesgenetics.org/gene/geneInfo/NQO1). NQO1 is highly expressed in human adipose tissue and its expression is reduced during diet-induced weight loss; furthermore its expression correlates directly with adiposity, glycaemia and markers of liver dysfunction[25]. Together, these findings indicate a role for NQO1 in the aetiology of obesity and type 2 diabetes.

**Inhibin Subunit Beta E (*INHBE*)** gene is a member of the Transforming Growth Factor (TGF) beta superfamily. The transcribed peptide Activin E is ubiquitously expressed in a large number of normal tissues, many being known to be especially active with cell proliferation, apoptosis, immune response and hormone secretion. The highest expression is found in the liver where it acts as a hepatokine with effects on energy homeostasis in both brown and white adipose tissue[26]. The candidacy of the *INHBE* gene is further supported by strong GWAS signals associating it with cardiometabolic traits, raised serum triglycerides and with coronary heart disease (http://www.type2diabetesgenetics.org/gene/geneInfo/INHBE).

### Others with suggestive evidence include

**Glutamate receptor interacting protein 1 (*GRIP1*)** mediates the trafficking and membrane organisation of a number of trans-membrane proteins in various cells including neurons and macrophages. Obese mice with a conditional knockout of *GRIP1* in macrophages develop massive macrophage infiltration and inflammation in many metabolically active tissues leading to many features that associate with the metabolic syndrome such as hepatic steatosis, hyperglycaemia and insulin resistance[27]. **Clusterin (*CLU*)** is a molecular chaperone. Secretory clusterin is also known as ApoJ. ApoJ has recently been identified as a novel hepatokine, and deletion of hepatic ApoJ leads to insulin resistance and glucose tolerance[28]. Furthermore, in humans, serum ApoJ levels correlate directly with increases in insulin resistance and these levels decrease in response to rosiglitazone treatment[29].

**Insulin Like Growth Factor Binding Protein 3 (*IGFBP3*)** is the most abundant of 6 IGF-binding proteins. Important interactions have been observed between IGFBP3, vitamin D metabolism and obesity[30]. Furthermore, in people with type 2 diabetes IGFB3 levels may inversely contribute to accelerated cerebrovascular disease[31]. **Potassium Voltage-Gated Channel Subfamily Q Member 5 (*KCNQ5*)** is a component of potassium channels. A strong GWAS association is seen between *KCNQ5* and body mass index (http://www.type2diabetesgenetics.org/gene/geneInfo/KCNQ5). **Cytotoxic And Regulatory T-Cell Molecule (*CRTAM***) is a Protein Coding gene that has a role in the innate immune system and has also been implicated as a potential determinant of insulin secretion[32]. A strong GWAS association is seen between *CRTAM* and both body mass index and systolic blood pressure (http://www.type2diabetesgenetics.org/gene/geneInfo/CRTAM). **Spectrin (*SPTA1*)** is a component of the erythrocyte plasma membrane. A strong association is seen between *SPTA1* and separately with HbA1c (GWAS) and type 2 diabetes adjusted for BMI (mainly missense mutations) (http://www.type2diabetesgenetics.org/gene/geneInfo/SPTA1).

A weakness of our approach is that although we have used a human immune cell line approach, we have not yet validated our findings *in vivo* in humans. Various compounds have been isolated and identified from *Areca catechu* nuts including alkaloids, tannins, flavones, triterpenes, steroids, and fatty acids. We have chosen to focus on arecoline and two of the several nitrosated products (MNPA and MNPN), selected as being the most carcinogenic of them, and in particular, because low-dose nitrosamines cause type 2 diabetes experimentally and in humans[12, 13]. Unfortunately, for technical reasons, we did not get results with MNPN. *Areca catechu* chewing quids often contain various other additives such as slaked lime, spices, sweeteners, and are wrapped in leaves of the *Piper betle* vine; furthermore, in many countries other than Taiwan they often contain chewing tobacco. We cannot therefore exclude the possibility that major effects of chewing betel quids in humans may be due to ingestion of betel quid components other than those from the *Areca catechu* nut. However, obesity and hyperglycaemia were induced in CD1 mice fed *Areca catechu* nut without any other betel quid component[9] and this data contributed to our focus on the findings for genes associated with those particular disorders in humans.

## Conclusion

This pilot study has identified a large number of genes whose expression changed significantly in human TPH1 cells following incubation with arecoline and MNPA and these genes are known to be associated with increased risks of obesity and type 2 diabetes in humans. These findings suggest that further investigation of these genes in betel-quid chewers with obesity and/or type 2 diabetes is warranted.

## Abbreviations

MNPA: 3-methylnitrosaminopropionaldehyde
MNPN: 3-methylnitrosaminopropionitrile
PMA: (Phorbol-12-Myrsitate-13-Acetate)

## Acknowledgements

The authors thank Nisha Patel (Borchukova laboratory, Blizard Institute, Queen Mary University of London) for technical support.

## Author contributions

SC, WO, CEM, ES performed the investigations and RL the bioinformatic analysis, GAH, WO and BJB supervised the study, GAH conceptualized the study and acquired the funding, GAH wrote the first draft of the manuscript with the help of BJB and all authors contributed to the writing of the manuscript thereafter. GAH is the guarantor of this work and, as such, had full access to all the data in the study and takes responsibility for the integrity of the data and the accuracy of the data analysis.

This article contains supplementary material

